# ChEA-KG: Human Transcription Factor Regulatory Network with a Knowledge Graph Interactive User Interface

**DOI:** 10.1101/2025.08.09.669505

**Authors:** Anna I. Byrd, John Erol Evangelista, Alexander Lachmann, Ho-Young Chung, Sherry L. Jenkins, Avi Ma’ayan

**Author notes:** Present Address: Avi Ma’ayan, Department of Pharmacological Sciences, Department of Artificial Intelligence and Human Health, Mount Sinai Center for Bioinformatics, One Gustave L. Levy Place, Box 1603, Icahn School of Medicine at Mount Sinai, New York, NY, 10029, USA.

## Abstract

Gene expression is controlled by transcription factors (TFs) that selectively bind and unbind to DNA to regulate mRNA expression of all human genes. TFs control the expression of other TFs, forming a complex gene regulatory network (GRN) with switches, feedback loops, and other regulatory motifs. Many experimental and computational methods have been developed to reconstruct the human intracellular GRN. Here we present a different approach. By submitting thousands of “up” and “down” gene sets from the RummaGEO resource for TF enrichment analysis with ChEA3, we distill signed and directed edges that connect human TFs to construct a high quality human GRN. The GRN has 131,581 signed and directed edges connecting 701 source TF nodes to 1,559 target TF nodes. The GRN is accessible via the ChEA-KG web server application, which provides interactive network visualization and analysis tools. Users may query the GRN for single or pairs of TFs or submit gene sets to perform TF enrichment analysis with ChEA3, placing the enriched TFs within the GRN. To demonstrate the utility of ChEA-KG, several TF-centric atlases are also made available via the ChEA-KG website. These atlases host TF subnetworks that regulate 131 major normal human cell-types (Cell Type Atlas); 69 tumour subtypes from 10 cancers (Cancer Atlas); 30 consensus perturbation response signatures for common mechanisms of action (MoA Atlas); and 24 aging signatures from tissues profiled by GTEx. Overall, ChEA-KG is an interactive web-server application that presents to users a new method of exploring the human gene regulatory network through both network visualization and transcription factor enrichment analysis. The ChEA-KG application is available from: https://chea-kg.maayanlab.cloud/.

## INTRODUCTION

Unravelling the regulatory mechanisms underlying genome-wide gene expression has been a major goal of experimental and computational biologists for decades. Transcription factors (TFs) are key controllers of gene expression. TF activity directly and indirectly responds to extracellular cues to determine the behaviour of the cell by binding and unbinding to specific DNA sequences to increase (upregulate) or decrease (downregulate) the production of transcripts. TFs regulate the expression of other TFs forming an intricate and complex regulatory network commonly termed gene regulatory network (GRN) [1, 2]. Understanding the connectivity and dynamics of GRNs is critical to explaining disease mechanisms and other key intracellular physiological processes. GRNs can be represented *in-silico* as directed and signed graphs with nodes representing TFs and edges representing their regulatory relationships. To infer these associations, experimental data that measure TF activity directly or indirectly can inform computational methods. For example, various experimental methods have been developed to identify the binding locations of TFs on the DNA. Such experimental techniques include chromatin immunoprecipitation followed by sequencing (ChIP-seq) [3] or microarray (ChIP-chip) [4]; open chromatin assays, including DNase sequencing (DNase-seq) [5], formaldehyde-assisted identification of regulatory elements followed by sequencing (FAIRE-seq) [6], and DNase footprinting [7]; and chromatin interaction assays such as chromatin interaction analysis with paired end tag sequencing (ChIA-PET) [8, 9]. Aggregating the results from many studies that utilized these assays can be used to reconstruct GRNs [10–13]. However, typically such GRNs only have a subset of the entire repertoire of TFs and contain directed but unsigned edges. Alternative and complementary computational methods to reconstruct GRNs include text-mining [14], position weight matrices (PWM) analysis [15], and TF-gene co-expression analysis [16]. However, these methods may suffer from literature biases, lack of specificity, and reliance on indirect evidence. Regardless of the advantages and disadvantages of the computational and experimental methods used to reconstruct GRNs, once these GRNs are formed, they can be analyzed for their topological features such as network motifs [17–19], hub analysis [20], positive and negative feedback loops [21] and other topological features [22, 23]. Moreover, a useful application of *in-silico* GRNs in biomedical research is their use as background knowledge for transcription factor enrichment analysis (TFEA) [10, 11, 24].

Several web server and command line tools are available for performing TFEA. For example, Genomic Regions Enrichment of Annotations Tool (GREAT) [24] takes as input genomics locations to predict enriched pathway and biological processes based on knowledge about the binding sites of TFs in the nearby regions. ChIP-x enrichment analysis (ChEA) [11] aggregates gene sets from ChIP-seq and ChIP-chip publications and serves these sets for standard enrichment analysis with a proportion test. The updated version of ChEA, ChEA3 [10] aggregates TF-target associations from other sources such as co-expression, protein-protein interactions, and PWMs to identify consensus enriched TFs. Other tools such as oPOSSUM [25], Pscan [26], HOMER [27], and i-cisTarget [28] rely only on PWMs to infer the most likely TFs given a set of genes or a file with genomics regions annotations. Several other tools developed to perform TFEA are available offering various features and input types (Table 1, Supplemental Table S1). Here, we introduce a different method to reconstruct the human TF GRN. By combining TFEA with ChEA3 [10] with a massive collection of differentially expressed gene sets created for establishing RummaGEO [29], we infer the most likely regulatory relationships between all human TFs. The inferred GRN is stored in a knowledge graph (KG) database and served via a web server application called ChEA-KG. ChEA-KG is an interactive TFEA application that returns subnetworks of the connected top enriched TFs based on the topology of the reconstructed GRN. ChEA-KG also provides interactive network visualization and allows querying for relationships within the full GRN. To demonstrate the utility of ChEA-KG for context-specific analyses, we created four atlases featuring curated collections of TF regulatory subnetworks predicted to control marker gene sets from 131 human cell types; subnetworks regulating gene expression in 69 cancer subtypes from 10 cancer types; subnetworks of TFs regulating changes induced by drugs with 30 major mechanism of actions (MoAs); and TF subnetwoks responsible for observed changes due to aging in 24 human tissues.

**Table 1.** Comparison the features of ChEA-KG with existing relevant published tools to predict TF regulatory interactions. A: Accessible in browser. B: Scripting library/package or requires download. C: Interactive visualization of results. D: Network of TFs. E: Infers sign of TF-target regulation. F: Background includes all human TFs.

| Name | PMID | Input | Output | A | B | C | D | E | F |
| --- | --- | --- | --- | --- | --- | --- | --- | --- | --- |
| **ChEA-KG** |  | Gene set | Subnetwork of enriched TFs and their connections | ✓ | × | ✓ | ✓ | ✓ | ✓ |
| **AME** | 25953851 | Genomic regions | Enriched motifs and TF binding sites | ✓ | × | × | × | × | × |
| **BART** | 29608647 | Gene set | Enrichment scores for TFs | ✓ | ✓ | × | × | × | × |
| **ChEA3** | 31114921 | Gene set | Ranked TFs predicted to regulate gene sets | ✓ | × | ✓ | × | × | × |
| **Cluster Profiler** | 22455463 | Gene set | Enrichment scores and visualization | × | ✓ | ✓ | ✓ | × | × |
| **DoRothEA** | 31340985 | Gene expression matrix | TF activity scores and regulatory networks | × | ✓ | ✓ | ✓ | ✓ | ✓ |
| **GREAT** | 20436461 | Genomic regions | Enriched terms across different gene set libraries | ✓ | × | × | × | × | × |
| **GTRD** | 33231677 | Genomic regions or gene lists | Enriched TF binding sites | ✓ | ✓ | ✓ | × | × | ✓ |
| **HOMER** | 20513432 | Genomic regions or gene lists | Enrichment of known or de novo motifs | × | ✓ | × | × | × | × |
| **iRegulon** | 25058159 | Gene set | Predicted TFs and their targets | × | ✓ | ✓ | ✓ | ✓ | ✓ |
| **PSCAN** | 19487240 | Genomic regions or gene lists | TF motif enrichment scores | ✓ | × | × | × | × | × |
| **RcisTarget** | 28991892 | Gene set | Ranked TFs associated with gene sets | × | ✓ | × | × | × | × |
| **TFEA** | 34079046 | Genomic regions or gene lists | Enrichment scores of TF motifs | × | ✓ | × | × | × | × |
| **TFEA.ChIP** | 31347689 | ChIP-seq peak files | TF enrichment scores | ✓ | ✓ | ✓ | × | × | × |
| **TFmotifView** | 32324215 | Genomic regions | Enriched TF motifs and their regulatory potential | ✓ | × | ✓ | × | × | × |
| **TRANSFAC** | 16381825 | Genomic regions or gene lists | Enriched TF binding sites and regulatory networks | ✓ | ✓ | ✓ | × | × | × |
| **TRRUST** | 26066708 | Gene set | TF-target networks with evidence from scientific literature | ✓ | × | ✓ | ✓ | ✓ | × |
| **Viper** | 27322546 | Gene expression signature and cell context-specific regulatory network | Inferred activity for each regulatory gene in the input dataset | × | ✓ | × | ✓ | ✓ | ✓ |

## METHODS

### Construction of the TF-TF GRN with TFEA

The GRN construction method assumes that the up- or down-regulation of genes resulting from a single perturbation are controlled by TFs that directly regulate the transcription of those genes. These upstream TFs can be identified by submitting many differentially expressed up-and down-regulated gene sets for TFEA with ChEA3 [10]. Differentially expressed gene sets were downloaded from RummaGEO [29] on August 22nd, 2024. Gene sets were kept if they had at least five genes. Only studies with clear control and perturbation groups were included to ensure that differential expression resulted in a clear direction of transcription of the target genes. These studies were identified using regular expressions for terms that describe the experimental conditions, for example, “ctrl” and “wt”; and by manual classification of samples by DiSignAtlas [30]. TFEA was performed on each gene set using a local version of ChEA3 [10]. The 10 top-ranked TFs for each gene set were identified using the MeanRank method, and source-target edges were counted between the top 10 enriched TFs (sources) and TF-encoding genes from the input set. The sign of the regulatory relationship was determined based on the direction of differential expression comparing the perturbation group to the control group (“up” or “down” gene sets). Edges with a count lower than 10 were discarded.

### Filtering the network to preserve significant edges

To remove edges that may be included by random chance, we determined the significance of each edge by comparing its observed frequency to its expected frequency in shuffled networks. Two methods were developed for generating expected edge counts. Both methods shuffle the original network. The “target set swap” (TSS) method swaps the source TF randomly, preserving the degree of the target nodes. The “node draw” (ND) method randomly swaps pairs of source and target nodes. Nodes are selected by the weighted frequency of occurrence as either a source or target role. This is repeated until the same number of edges have been sampled as were counted in the unfiltered network, preserving the relative connectivity distribution of the source and target nodes. For each shuffling method, 50 shuffled networks are produced. Then, the average and standard deviation of each edge is calculated. Next, the observed edge counts are compared to the average and standard deviations of the counts of each set of shuffled networks to calculate a z-score for the edge. The z-score was used to calculate a right-tailed p-value. Edges with a p-value >= 0.01 were discarded. Due to the large number of initial edges, some source-target node pairs were connected by both upregulated and downregulated edges. This conflict was resolved by preserving only the sign of the more significant edge. This produced two filtered GRNs: TSS-filtered and ND-filtered. Each of these GRNs was evaluated for accuracy by comparing the edges within these networks to edges from other sources of TF-TF associations not used to construct the GRNs.

### Benchmarking the filtered networks against other GRNs

Each filtered network was independently benchmarked against two reference GRNs to determine if the GRN reconstruction process produces known TF-target associations, as well as which network pruning method produces the most reliable network. The two reference GRNs used to benchmark the ChEA-KG GRN are from text mining of TF-target interactions from TRRUST [14], and position weight matrices (PWMs) from TRANSFAC [31] and JASPAR [15]. These networks were selected because they are not used by ChEA3 [10] to perform enrichment analysis and were processed and available for download from Enrichr [32]. The TRRUST GRN has 379 source TFs, 553 target TFs, and 2,597 edges. The TRANSFAC and JASPAR PWMs GRN contains 246 source TFs, 1,652 target TFs, and 32,067 edges. The statistical properties of the ChEA-KG filtered and unfiltered networks are summarized in Supplementary Table S2. The filtered and unfiltered ChEA-KG GRNs were compared to the benchmarking GRNs to determine the number of overlapping edges, and whether the number of overlapping edges is greater than what would be expected in shuffled networks with the similar size and structure. To generate the shuffled networks, source-target pairs in the original ChEA-KG filtered and unfiltered networks were randomly sampled from the source and target node distributions. This process ensures that each shuffled network has the same number of nodes and edges and similar connectivity distribution. The number of overlapping edges were normalized to the size of the filtered network. To determine statistical significance of the expected vs. observed overlapping edges between the ChEA-KG original and shuffled networks the Z test was applied. 100 shuffled networks were generated to produce an expected overlap distribution. The benchmarking procedure is outlined in a diagram (Supplementary Figure 1). This procedure was performed twice for each comparison, once with directed edges, and once with undirected edges, for a total of eight comparisons.

### Visualizing the GRN

To better understand the global structure of the GRN, several dimensionality reduction methods were applied to visualize and cluster the GRN’s TFs based on their connectivity similarity. First, we created a uniform manifold approximation projection (UMAP) [33] of the source TFs based on similarity between their target TF sets. Term frequency-inverse document frequency (TF-IDF) for each TF were computed [34]. Then, the Leiden algorithm was applied to identify clusters of similar TFs [35]. Next, we visualized the network edges as a clustered heatmap. To create the heatmap, first the network was converted to a directed, signed adjacency matrix where rows represent sources, and columns represent targets. Each (source, target) entry in the matrix was assigned a 0 if no edge exists, or a 1 or −1 for upregulated or downregulated edges, respectively. In addition, a heatmap of the Jaccard similarity scores between all TF pairs was generated. We calculated the Jaccard similarity between each pair of source TFs based on their shared targets. Both matrices were plotted as hierarchically clustered heatmaps using the Seaborn Python package [36].

### Developing the ChEA-KG web server application

The ChEA-KG web-server application enables users to interact with and query the GRN. The application was built using the customizable Knowledge Graph User Interface (KG-UI) that uses Cytoscape.js [37] to visualize Cypher query results from a Neo4j database [38]. The GRN was ingested into a Neo4j database by serializing the GRN into node and edge lists. The user interface (UI) provides the ability to perform queries for finding neighbours of single TFs, finding shortest paths between pairs of TFs, displaying the subnetworks returned from the enrichment analysis with ChEA3 [10], displaying the Cell Atlas subnetworks using various layouts, expanding and shrinking the size of the displayed subnetwork, viewing properties of nodes and links, and downloading the displayed associations in tabular format.

### Building the TF Cell Atlas

Cell types for the TF subnetwork Cell Atlas were identified manually based on literature curation. Blood cell types appearing in multiple tissues were removed. To construct the subnetworks for each cell type, cell-type specific marker gene sets were extracted from CellMarker [39], ASCT+B [40], PanglaoDB [41], Tabula Sapiens [42], Descartes [43], Azimuth [40], and TISSUES [44]. These marker genes were identified by manually mapping cell types to the terms that represent gene sets in gene set libraries served by Enrichr [32]. One gene set was selected for each cell type. Once the gene sets were present for each cell type, the gene sets were submitted to ChEA3 [10] for enrichment analysis and the networks were plotted using the KG-UI. We further analysed six representative subnetworks within the Cell Atlas to identify literature evidence linking each cell type to the nodes and link in each subnetwork of inferred TFs. Relevant citations were identified by manually searching PubMed for the cell type, tissue type, and TF. For TFs with common synonyms documented in the NCBI database, for example, SPI1 and PU.1, both names were considered in the search. Relationships between the TFs within the subnetworks, including structural similarity, co-expression, known physical interactions, and transcription regulation were recorded.

### Building the TF Cancer Atlas and analysing the subnetworks with GSFM

The ChEA-KG Cancer Atlas was created by submitting to ChEA-KG gene sets that are identified as upregulated in the transcriptomics clusters of the 10 tumour types from the Clinical Proteomic Tumour Atlas Consortium (CPTAC3) cohort [45]. For each cluster, identified subtypes were detected with the Leiden clustering algorithm [35] as described in the Multiomics2Targets workflow [46]. The upregulated marker gene sets for each subtype were extracted from the output of Multiomics2Targets, which uses the limma-voom to compute differential expression [47]. The visualization and integration of the ChEA-KG Cancer Atlas follows the same steps described for setting up the ChEA-KG Cell Atlas. To analyse the collective functions of each subnetwork produced by ChEA-KG, we identified consensus function predictions using the Gene Set Foundation Model (GSFM) [48] the KOMP2 [49] Mouse Phenotypes 2022 library from Enrichr [32] for the TFs in each subnetwork. The top 10 predictions with the highest score for each subnetwork of TF were aggregated and then sorted based on z-score. The top 10 aggregated phenotypes were assigned as the consensus phenotype for each subnetwork.

### Building the TF MoA Atlas

The ChEA-KG MoA Atlas was created using precomputed differentially expressed gene sets from the LINCS L1000 Fireworks Display (L1000FWD) application [50]. Gene sets were downloaded directly from the L1000FWD download page. The set of unique drugs were identified and then cross referenced with the Drug Repurposing Hub [51] to assign known MoAs to each drug. All drugs with no known MoA were labelled as “unknown” and were discarded. The top 30 most common MoAs were selected as the set of terms for the gene set library. Up-regulated and down-regulated gene sets were produced for each MoA for a total of 60 gene sets. To create the consensus gene sets for each term, first, all the up- or all down-regulated genes for each MoA drug were counted. A consensus cutoff was then iteratively reduced until both up and down gene set size averages were between 150 and 200 genes. To quantify the overlap between each gene set, the Fisher’s exact test. Three comparisons were performed: upregulated vs. upregulated, upregulated vs. downregulated, and downregulated vs. downregulated gene sets. The p-values were recorded for each comparison and visualized as hierarchically clustered heatmaps using the Seaborn Python package [36]. The visualization and integration of the MoA Atlas follows the same procedure as the previously described Atlases. To identify shared modules of TFs between different MoAs, an adjacency matrix representing all edges across the 60 subnetworks was constructed. First, unique edges across all networks were combined into a single network. A directed binary adjacency matrix was created where 1 indicates a source-target edge and 0 indicates no edge. The resulting binary matrix was visualized as a hierarchically clustered heatmap using the Seaborn Python package [36]. Modules of similar TFs were identified and extracted. Their corresponding subnetworks were identified through manual inspection of the heatmap.

### Building the TF Aging Atlas

The ChEA-KG Aging Atlas was created using differentially expressed gene sets pre-computed for the CFDE Gene Set Cart project [52] using the GTEx [53] Aging Signatures dataset. The 20-29 vs. 70-79 age group comparison was used as the representative set of signatures, up/down gene sets, for each tissue. The visualization and integration of the Aging Atlas follows the same procedure as the described for the other Atlases. The resulting subnetworks were tested for significant overlap to identify tissues with similar TF subnetworks. Significance was assigned via permutation testing. Given a pair of subnetworks A and B with sizes M and N, respectively, and overlapping edges S_0_, 1000 pairs of networks with sizes M and N and overlap S_1_ were generated by randomly sampling edges from the full ChEA-KG background GRN. The p-value was calculated by dividing the number of times the randomized overlap, S_0_, is greater than or equal to the observed overlap, S_1_, divided by the number of permutations (1000). A pan-tissue regulatory TF-TF signed and directed network was constructed by aggregating all edges present in the Aging Atlas subnetworks. The resulting global network was visualized using Cytoscape [37]. The frequency of observing each node and link in the Aging Atlas subnetworks informs the size of the nodes and links in the global network.

## RESULTS

### The TF-TF GRN

The ChEA-KG human GRN of TF-TF interactions was created by applying TFEA with ChEA3 [10] to a massive collection of gene sets created by automated differential gene expression analysis applied to thousands of RNA-seq studies deposited in GEO to create RummaGEO [29]. By combining TF binding site knowledge with differential gene expression data, we can capture the most likely direct effects of one TF over another to construct a high fidelity GRN (Figure 1). The RummaGEO resource serves 171,441 human and 195,265 mouse gene sets spanning 29,294 GEO studies. For establishing ChEA-KG, only a subset of 29,328 gene sets extracted from 10,901 unique studies were used because the sets from these studies came from signatures where there is a clear separation between a control and a perturbation group. These gene sets were created by first downloading the uniformly aligned sample vectors of expression from ARCHS4 [54]. Then, samples from studies with control and perturbation conditions were analysed for differential expression with the limma-voom method [47]. TFEA was performed using ChEA3 [10] which covers 1,632 human TFs. During the construction process 10,405,833 signed edges were established between TFs. 373,250 of these edges are unique, connecting 727 source TFs to regulate the expression 1,560 target TFs. The network has self-loops and TFs can serve as both sources and targets. The GRN was filtered to exclude edges that did not occur more than expected when comparing the GRN to shuffled networks. After filtering, the GRN was reduced to 131,181 edges (ND filtering method) and 109,813 edges (TSS filtering method). The connectivity distributions of each of the filtered networks and the unfiltered network are visually compared (Figure 2). Other network properties such as number of edges by direction, average links per node, self-loops, and two-node feedback loops were recorded (Table 2).

**Figure 1.**
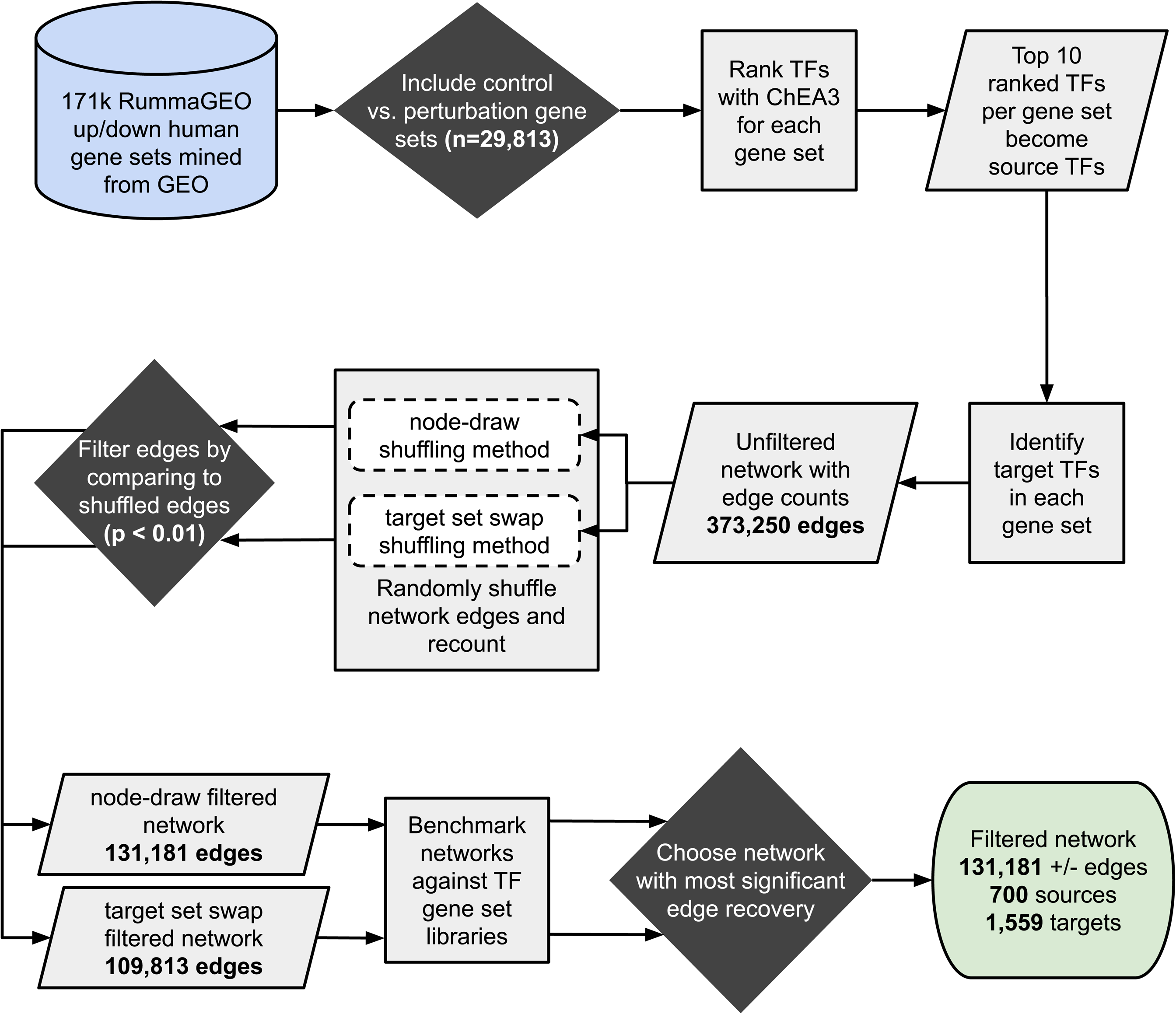
Workflow for constructing the GRN.

**Figure 2.**
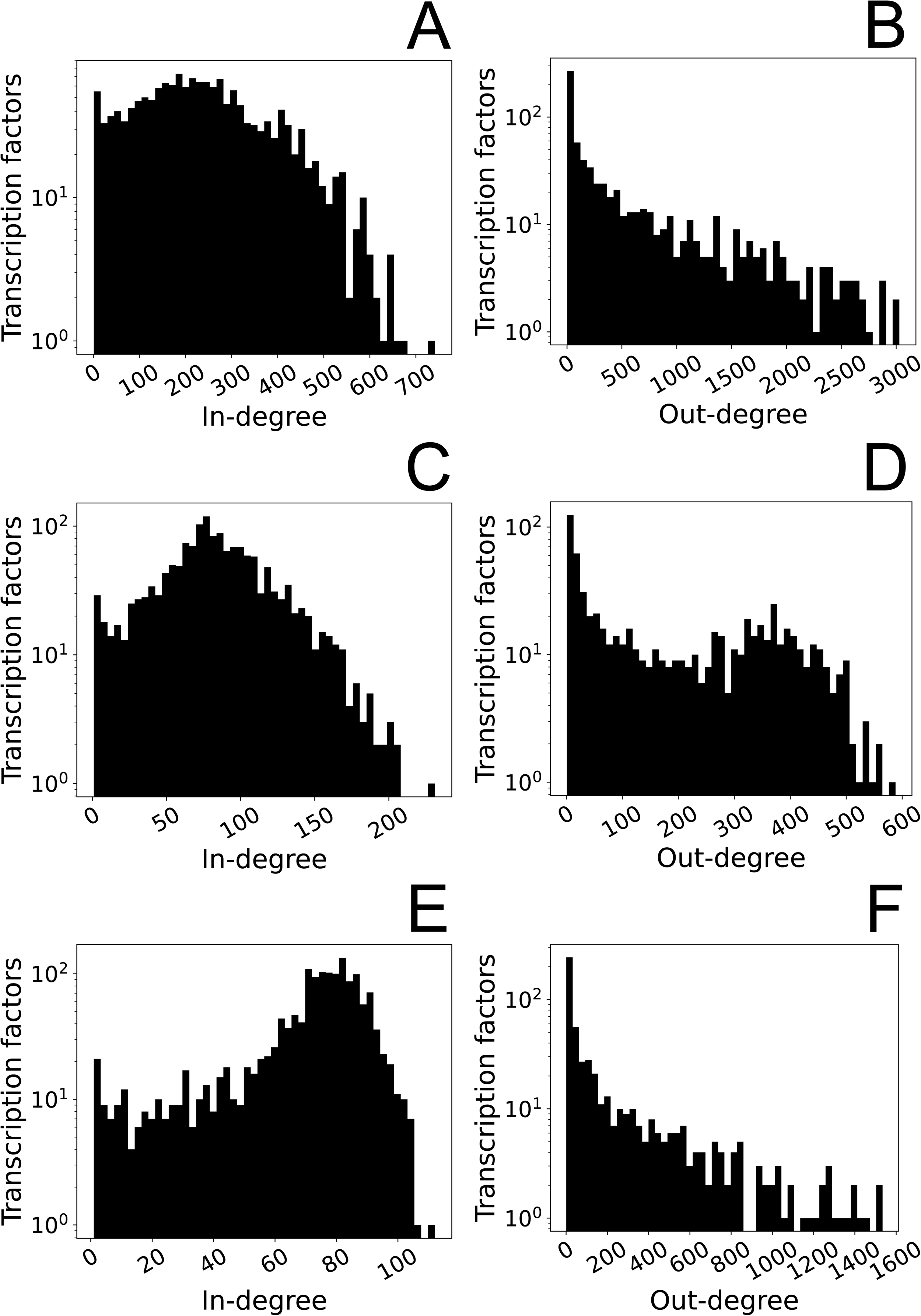
Network connectivity distributions of the unfiltered and ND- and TSS-filtered GRNs. In-degree and out-degree refer to target and source nodes, respectively. A, B: before filtering, C, D: ND-filtered; E, F: TSS-filtered.

**Table 2.** Network statistics. The nodes, edges, links per node, and self- and two-node feedback loops are of the unfiltered GRN and ND- and TSS-filtered GRNs. Edges in positive feedback loops have the same sign. Edges in negative feedback loops have opposite signs.

|  | Unfiltered network | ND-filtered | TSS-filtered |
| --- | --- | --- | --- |
| **Nodes** | 1,560 | 1,559 | 1,560 |
| **Source nodes** | 727 | 700 | 548 |
| **Target nodes** | 1,560 | 1,559 | 1,560 |
| **Edges** | 373,250 | 131,181 | 109,813 |
| **Upregulated edges** | 184,134 | 63,531 | 51,382 |
| **Downregulated edges** | 189,116 | 67,650 | 58,431 |
| **Out-links per node** | 239.263 | 84.144 | 70.393 |
| **In-links per node** | 513.411 | 187.401 | 200.389 |
| **Links per node** | 239.263 | 84.144 | 70.393 |
| **Self-loops** | 1030 | 549 | 379 |
| **Positive feedback loops** | 27,368 | 5,088 | 2,243 |
| **Negative feedback loops** | 27,185 | 4,514 | 2,075 |

### Benchmarking the ND and TSS filtering methods

Since the TSS- and ND-filtered GRNs have different nodes and edges, the two methods were benchmarked by comparing the edges in those networks to edges from other TF-TF GRNs that are not included in ChEA3 [10] and RummaGEO [29], namely, literature curated TF-TF interactions from TRRUST [14], and TF-TF interactions based on position weight matrices (PWMs) from TRANSFAC [31] and JASPAR [15]. Each comparison was made with and without considering edge direction, for a total of four comparisons per network (Figure 3). Both filtered GRNs showed significantly higher overlap with interactions from TRRUST and TRANSFAC/JASPAR compared to edges in the shuffled networks of the same size and similar connectivity. The ND-filtered GRN showed the most significant results in three out of the four comparisons. It was only outperformed by the TSS-filtered GRN when compared against the TRANSFAC/JASPAR PWMs library (Figure 3, Supplementary Table S2). Based on the benchmarking result the ND-filtered GRN was selected for the ChEA-KG webserver application.

**Figure 3.**
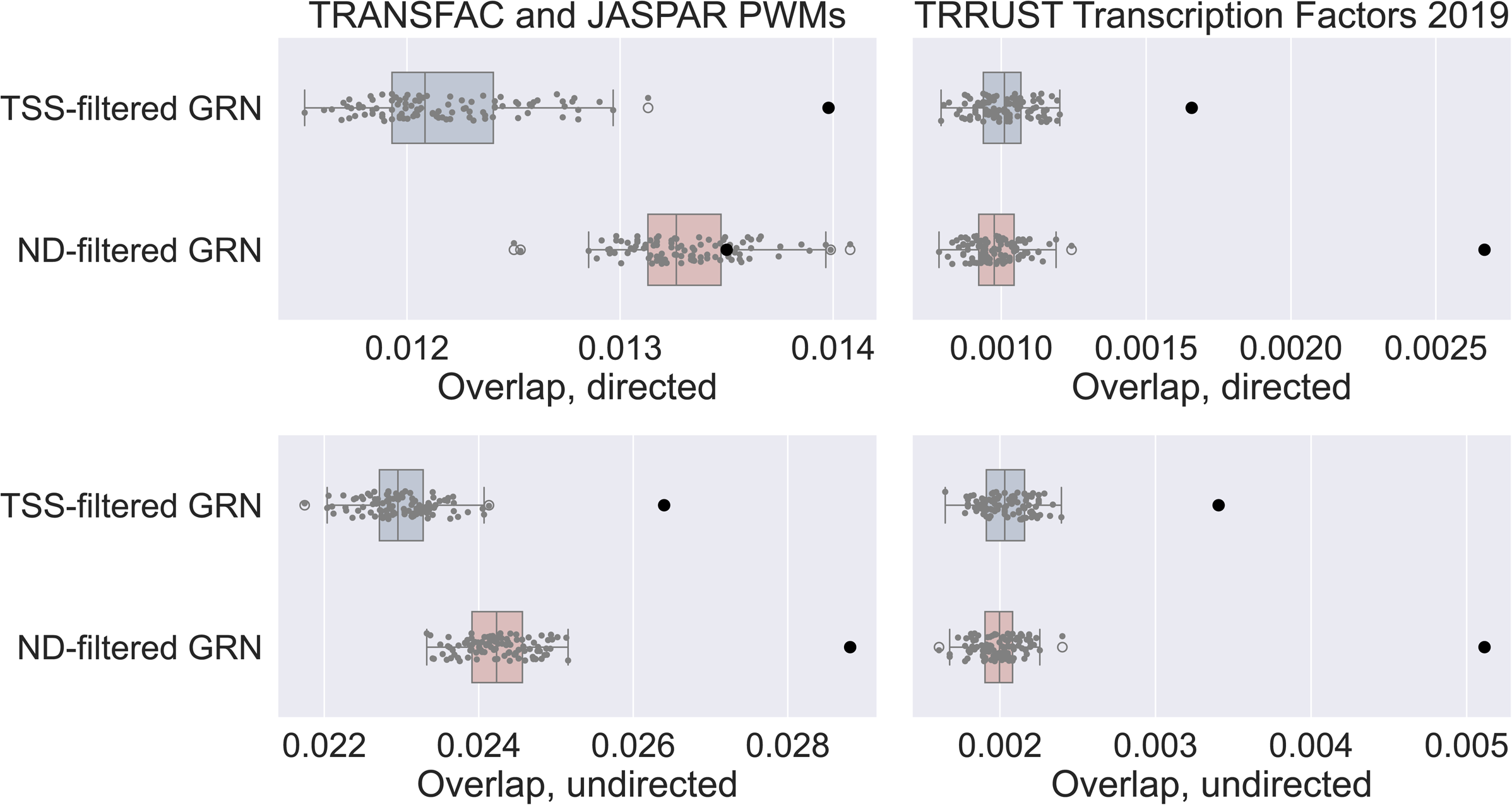
Benchmarking edge pruning methods (filtering). The expected edge overlap, calculated using randomly shuffled networks (shown with boxplots), is compared to the observed edges in the GRN (black dot). Comparisons are made with and without considering edge direction. The boxplots show the distribution of random trials. The black dots are the observed recovery for each comparison.

### Identifying clusters within the GRN

Next, we visualised the entire GRN to examine its topology and identify TF modules. First, the network was clustered with the Leiden algorithm [35] and then visualized with a UMAP [33] (Figure 4). The sets of TFs from the 26 identified clusters were uploaded to Enrichr [32] to characterize the collective functions of each cluster of TFs. 13 clusters had clearly defined collective functions such as “patterning”, “cell cycle”, “hair and skin regulation”, and “stem cell maintenance”. Next, the GRN adjacency matrix was visualized as a clustered heatmap (Supplementary Figure S2). Only the 700 TFs that act as both sources and targets are included in the heatmap. The densely coloured areas in the heatmap show several regions of TF modules that regulate similar targets. Lastly, the Jaccard similarity scores between GRN TFs were visualized as a clustered distance table heatmap (Figure 5). Only 381 highly connected TFs were visualised because this subset of TFs had clearly visible defined clusters. The dendrogram from the distance table was extracted, and clusters were determined by cutting the dendrogram at 65% of the maximum tree height to produce 12 unique clusters. These clusters were annotated with enriched terms identified with Enrichr. The heatmap clusters were enriched for terms such as “protein folding”, “immune response”, and “cell cycle”.

**Figure 4.**
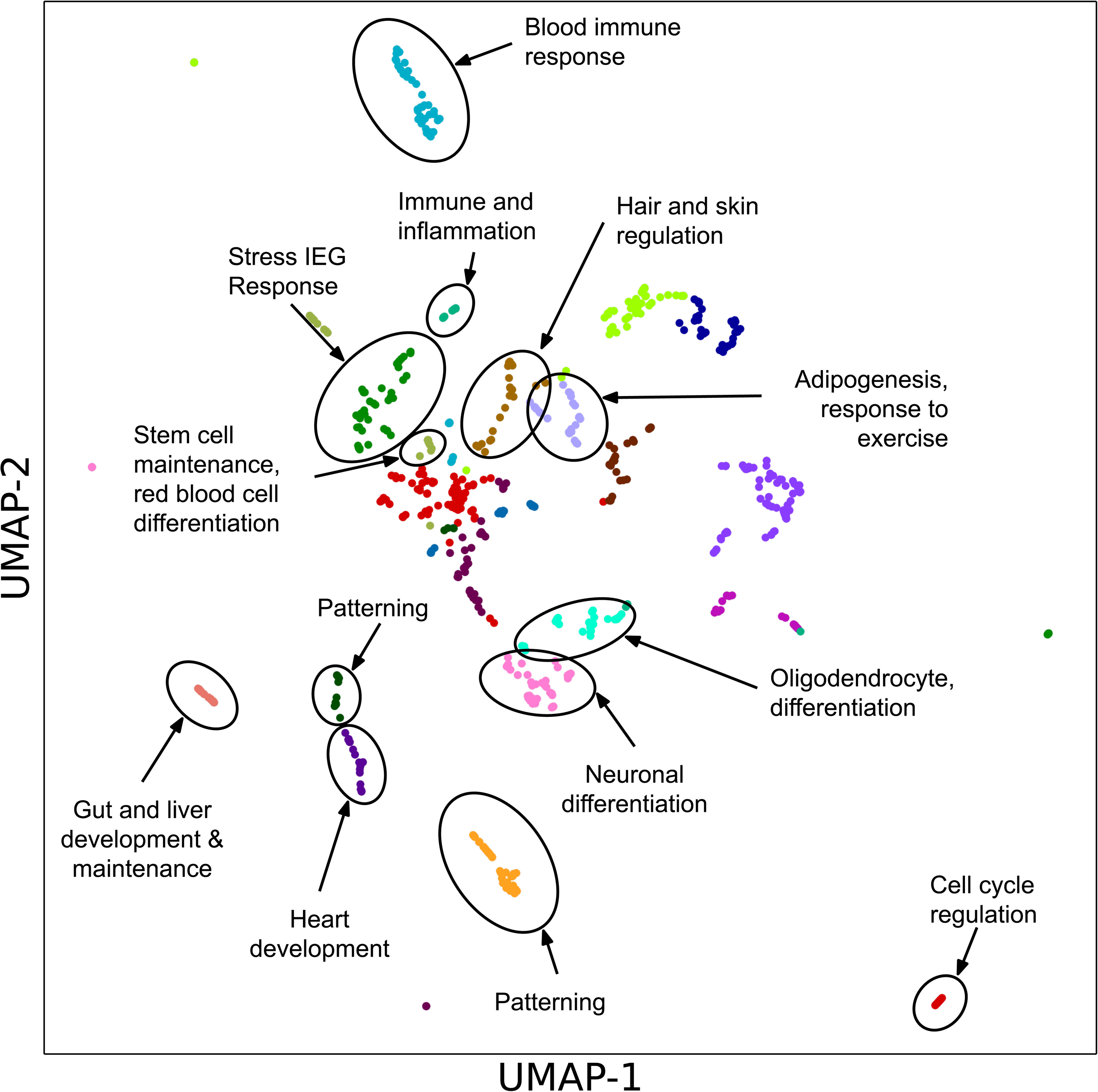
UMAP visualization of source TFs based on their target set similarity. Clusters are annotated with enriched terms.

**Figure 5.**
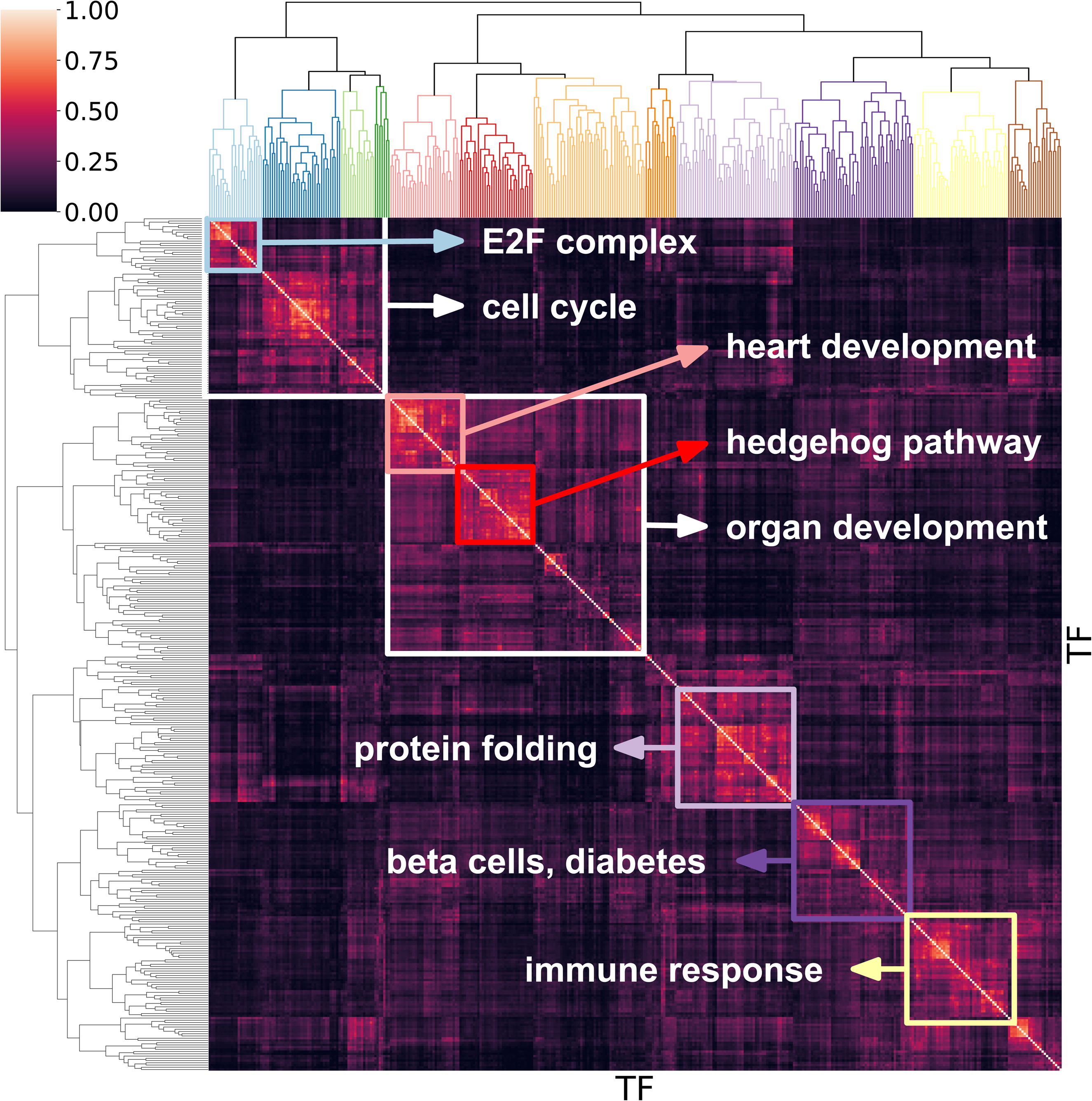
Clustermap of TFs based on their Jaccard similarity score computed based on their edge similarity. The dendrogram at the top is color-coded by cluster. Regions of high similarity are annotated with their enrichment terms.

### The ChEA-KG web server application

ChEA-KG is an interactive web server application that enables users to interact with the GRN in multiple ways. The human GRN is stored in a Neo4j database as a KG and the ChEA-KG UI is querying the KG database to extract and display GRN subnetworks. Users of ChEA-KG can query the KG database for creating subnetworks centred on a single TF, and subnetworks that connect two-TFs based on the shortest paths between them. The results page has a toolbar that enables users to customize the subnetworks by limiting the number of nodes and edges, viewing the subnetwork as a table, adding a legend, changing the subnetwork layout, and displaying a tooltip with information associated with each TF. Users can also save the subnetwork as an image, or as node and edge lists in CSV files. ChEA-KG also has an enrichment analysis feature that enables users to associate their input gene sets with subnetworks of regulatory interacting TFs. The input gene sets are queried using the ChEA3 API [10] to identify the top enriched TFs for each gene set. These TFs are then visualized as a subnetwork. Users can adjust the number of edges in the displayed subnetwork by filtering by significance. The number of TFs can be adjusted to return between 5-25 top enriched TFs. Because the number of TFs in the GRN (1,559) is less than the number of TFs in the ChEA3 database (1,632), it is possible that some top-ranked TFs are not connected to any other TF in the GRN. Even for TFs that are in the GRN, the enrichment results may return TFs that are not directly connected to each other. If an enriched TF does not share edges with any other enriched TFs, including self-loops, they will be displayed as a single unconnected node.

### The TF subnetworks cell atlas

Another feature of ChEA-KG is the TF subnetworks Cell Atlas. To create the Cell Atlas, we manually identified 131 major human cell types that can be found in 14 major human tissues. We then identified marker gene sets associated with each cell type by processing data from CellMarker [39], ASCT+B [40], PanglaoDB [41], Tabula Sapiens [42], Descartes [43], Azimuth [40], and TISSUES [44]. Then, we submitted these marker gene sets for enrichment analysis with ChEA3 [10] to create a subnetwork for each cell type. The TF Cell Atlas subnetworks are available through the ChEA-KG application. Users can select a subnetwork to view and interact with by using a drop-down menu of cell types organized by tissue. The full listing of cell types and marker genes are organized into a gene matrix transpose (GMT) file (Supplementary Table S3). From the collection of 131 cell types, we investigated the literature evidence of six subnetworks: erythrocytes in blood, cardiac muscle cells in heart, goblet cells in the intestine, mesenchymal stromal cells (MSCs) in bone marrow, epithelial cells in lung, and podocytes in the kidney (Figure 6).

**Figure 6.**
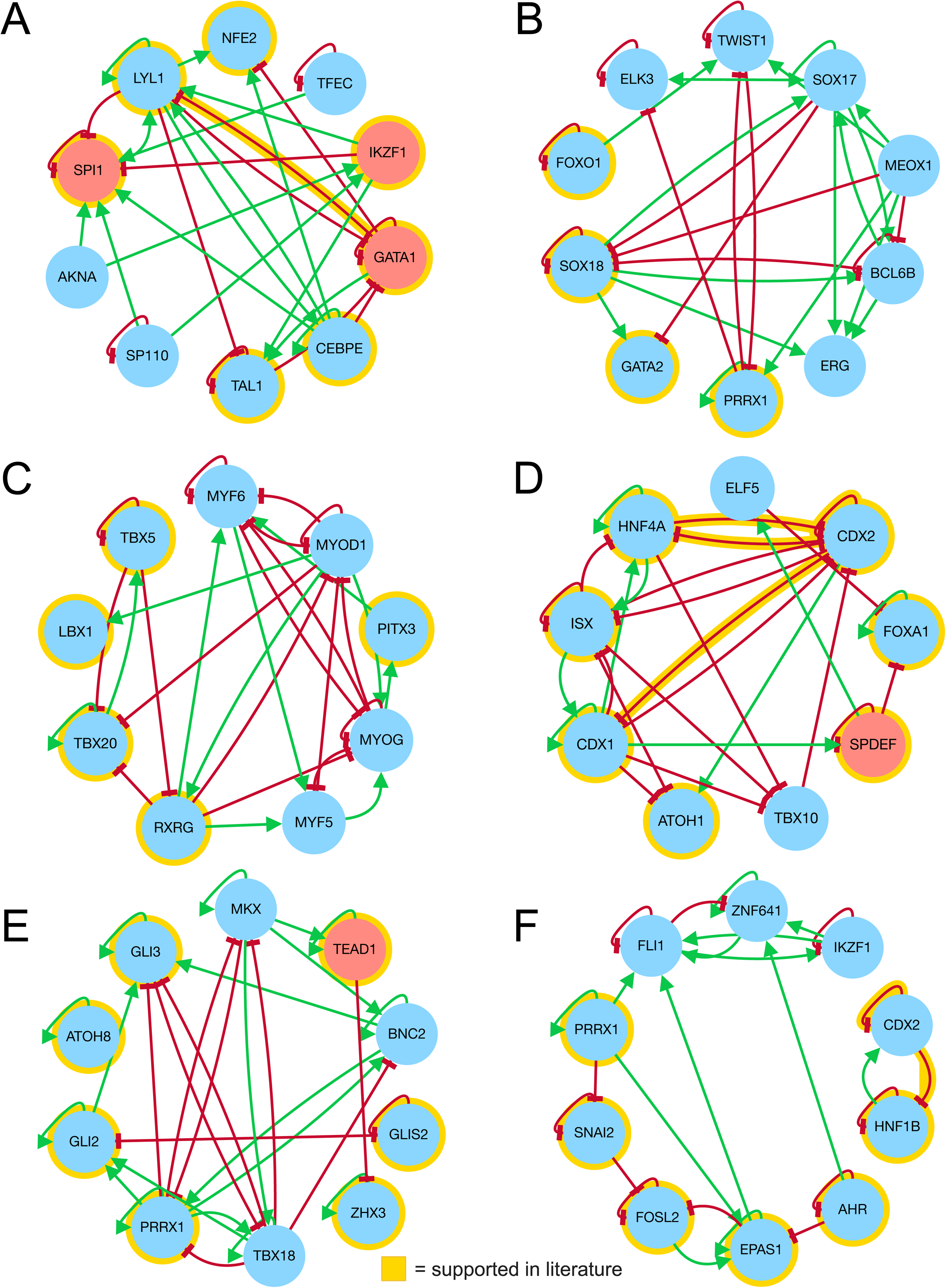
Six selected cell type specific TF regulatory subnetworks from the ChEA-KG TF Cell Atlas. A: Erythrocytes (blood). B: Mesenchymal stem cells (bone marrow). C: Cardiac muscle cells (heart). D: Goblet cells (intestine). E: Podocytes (kidney). F: Epithelial cells (lung). Edges and nodes are highlighted in yellow if they are confirmed by a manual literature search.

#### Blood: Erythrocytes

TFs and interactions in the erythrocyte regulatory subnetwork were found to be involved in several erythrocyte-specific processes. NFE2 is known to control the expression of globin genes in erythrocytes [55]; TAL1 [56], GATA1 [57], IKZF1 [58], SPI1 [59], and LYL1 [60] are known regulators of erythrocyte differentiation. TFs within the subnetwork are directly connected via direct transcriptional regulation, physical interactions, and structural similarity. LYL1 is known to bind to GATA1 to maintain erythropoiesis [61]. GATA1 also physically interacts with SPI1 [62] and controls the transcription of TAL1 [63], an interaction that is supported by the subnetwork (Figure 6A). The TFs AKNA, CEBPE, and TFEC were not found to be directly associated with erythrocytes in the literature and thus may represent undiscovered regulators of erythropoiesis.

#### Intestine: Goblet cells

The goblet cell regulatory subnetwork features several TFs known to control development of the intestine. CDX2, CDX1, and ATOH1 are known to regulate intestinal epithelial development [64], whereas HNF4A, SPDEF, and FOXA1 specifically regulate goblet cells by controlling maturation and differentiation [65–67]. ELF5 is a regulator of epithelial cell differentiation but has not been specifically linked to the intestine [68]. The T-Box family of genes plays a significant role in early embryonic development, but the exact role of TBX10 is unknown [69]. SPDEF and ATOH1 are transcriptional co-regulators [70] (Figure 6D). Overall, this subnetwork contains some known and some unknown master regulators of goblet cells.

#### Heart: Cardiac muscle cells

The cardiac muscle cell subnetwork includes TFs associated with both cardiac and skeletal muscle cells. RXRG2, an isoform of RXRG, is known to be highly expressed in cardiac muscle [71]. Several TFs in the cardiac muscle cell subnetwork are known regulators of skeletal muscle. For example, PITX3 is involved in myogenesis and has sustained expression in skeletal muscle cells [72]. MYF5, MYF6, MYOG, and MYOD1 are well-documented to establish the skeletal muscle phenotype [73]. These four genes share structural similarity and are collectively known as the Myogenic regulatory factors (MRFs) [74]. TBX20 and TBX5 are T-box proteins that are required for cardiomyocyte proliferation and homeostasis [75] and maturation [76], respectively. LBX1 controls gene expression relating to migration of muscle precursors [77]. Within the subnetwork, the skeletal muscle TFs, the MRFs and PITX3, regulate each other but no other network nodes, except for MYOD1, which regulates both RXRG and TBX20 (Figure 6C).

#### Kidney: Podocytes

TFs involved in the podocyte subnetwork are related to differentiation and gene expression during injury (Figure 6E). ATOH8 is a known regulator of kidney development and podocyte differentiation [78]. Several TFs in the subnetwork are associated with disease or injury of the kidney. ZHX3 is known to regulate podocyte gene expression during kidney injury [79, 80]. TEAD1 expression is lowered in diabetic podocyte injury [80]. In the kidneys more broadly, GLI2 is upregulated after adriamycin neuropathy [81] whereas GLIS2 is known to maintain renal architecture [82] and GLIS3 is associated with renal anomalies [83]. Finally, PRRX1 is suppressed during regulation of the renal epithelial-to-mesenchymal transition (EMT) [84], a process linked to podocyte dysfunction [85].

#### Lung: Epithelial cells

The epithelial cell subnetwork contains TFs involved in mucin production as well as EMT and lung cancer (Figure 6F). The TF SNAI2 plays a central role in EMT [86]. PRRX1 is similarly known to induce EMT in embryonic stem and cancer cells [87]. AHR is known to upregulates the expression of mucin 5AC [88], a major component of mucus in the lung [89]. High expression of FOSL2 has been linked to poor prognosis in lung cancer patients [90]. Lastly, HNF1B regulates the expression of alpha(1)-antitrypsin (AAT; SERPINA1), which may defend the lung against antiproteases [91]. CDX2, which is similarly enriched in intestinal epithelial cells, autoregulates its own expression and as well as the expression of HNF1B within intestinal epithelial cells, suggesting that it plays a similar regulatory role in lung and intestinal epithelial cells [92].

#### Bone marrow: MSCs

TFs enriched for MSCs are known to be involved in bone development and maintenance of processes related to cell fate (Figure 6B). SOX18 is known to be involved in the differentiation of MSCs to endothelial cells [93]. SOX17 is known to control the expression of fetal hematopoietic poetic stem cells [94], whereas MEOX1 is a key regulator of proliferation in mesenchymal-like cells [95]. Differentiation of bone-marrow-derived MSCs is regulated by GATA2 [96]. ERG and ELK3 are both members of the ETS family of TFs [97] and involved in EMT [98]. ERG is directly linked to endo-cardial mesenchymal transformation (EnMT) and EMT [99, 100]. Finally, it is known that TWIST1 suppresses the expression of RUNX2 to suppress bone formation [101] while FOXO1 defends against oxidative damage in the bone [102]. In conclusion, the subnetworks for each cell type in the ChEA-KG Cell Atlas confirm many master regulators as well as predict novel key regulators.

### The TF subnetworks cancer atlas

Besides the ChEA-KG Cell Atlas, ChEA-KG also has a Cancer Atlas. To create the ChEA-KG Cancer Atlas we obtained 69 up-regulated gene sets extracted from 10 cancer types profiled by the National Institutes of Health (NIH) CPTAC3 program [45]. Each of these gene sets represents a subtype based on applying the Leiden clustering algorithm [35] to the RNA-seq profiling of over a thousand tumours by Multiomics2Targets [46]. These up-regulated genes from these tumour subtypes were submitted for analysis with ChEA-KG. As an example, we applied this analysis to identify TF subnetworks enriched for lung squamous cell carcinoma (LSCC) (Figure 7). To characterize these subnetworks, we next identified key regulatory mechanisms for each LSCC subnetwork by predicting gene functions for each TF in each subnetwork using GSFM [48] and the KOMP2 [49] Mouse Phenotypes 2022 gene set library. (Supplementary Table S5). We find that cluster 0 is enriched in functions related to T-cell regulation, cluster 1 is enriched for midbrain morphology and sclerocornea phenotypes, cluster 2 is shows enriched terms for immunity and NK cells, cluster 3 appears to be related to lung development, and cluster 5 is related to thyroid regulation. These predicted functional terms immediately suggest key mechanisms of each cancer subtype, and for classifying tumours as hot or cold. The analysis provides clear mechanisms that may be correlated with subtype survival and likelihood of the responsiveness to therapy for each subtype.

**Figure 7.**
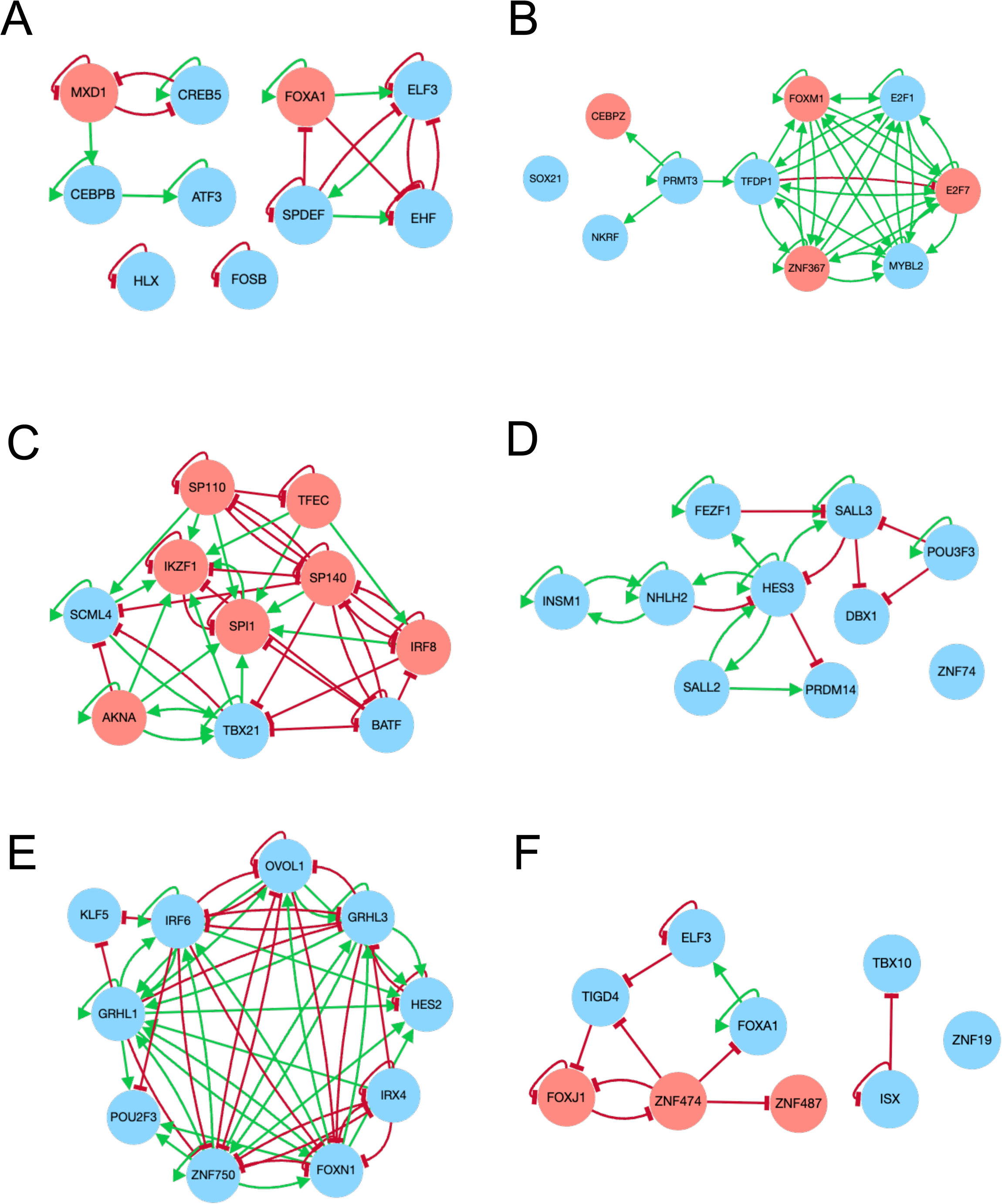
TF regulatory subnetworks identified by ChEA-KG from the six transcriptomics subclusters of LSCC from the ChEA-KG TF Cancer Atlas. A: Cluster 0 (n=29 patients/tumors). B: Cluster 1 (n=27). C: Cluster 2 (n=20). D: Cluster 3 (n=16). E: Cluster 4 (n=14). F: Cluster 5 (n=8). Red nodes represent TFs that are enriched by ChEA3 but also present in the input set of upregulated genes in the specific cluster.

### The TF subnetworks MoA atlas

The third ChEA-KG atlas is the MoA atlas. To create this atlas, we identified the 30 most common MoAs in the LINCS L1000 chemical perturbations dataset [103]. We then created consensus up and down gene sets for each MoA by aggregating genes that are up- or down-regulated in at least 17% of the signatures associated with each MoA (Supplementary Table S6). The gene sets were analysed for significant overlap using Fisher’s exact test (Supplementary Figure S3). A summary of each gene set, including the number of signatures and unique drugs used to compute the gene set was computed (Table 3). To identify key TF modules involved in drug response, we aggregated all unique edges from the MoA atlas subnetworks and generated a hierarchically clustered binary adjacency matrix heatmap. We identified five unique clusters of TFs with similar regulatory patterns (Figure 8A). For each cluster, we used the member TFs to assign one or two representative subnetworks from the MoA atlas (Figure 8B-F). We investigated the literature evidence connecting the TFs in clusters 1, 2, 4, and 5 to each of their MoAs as follows.

**Figure 8.**
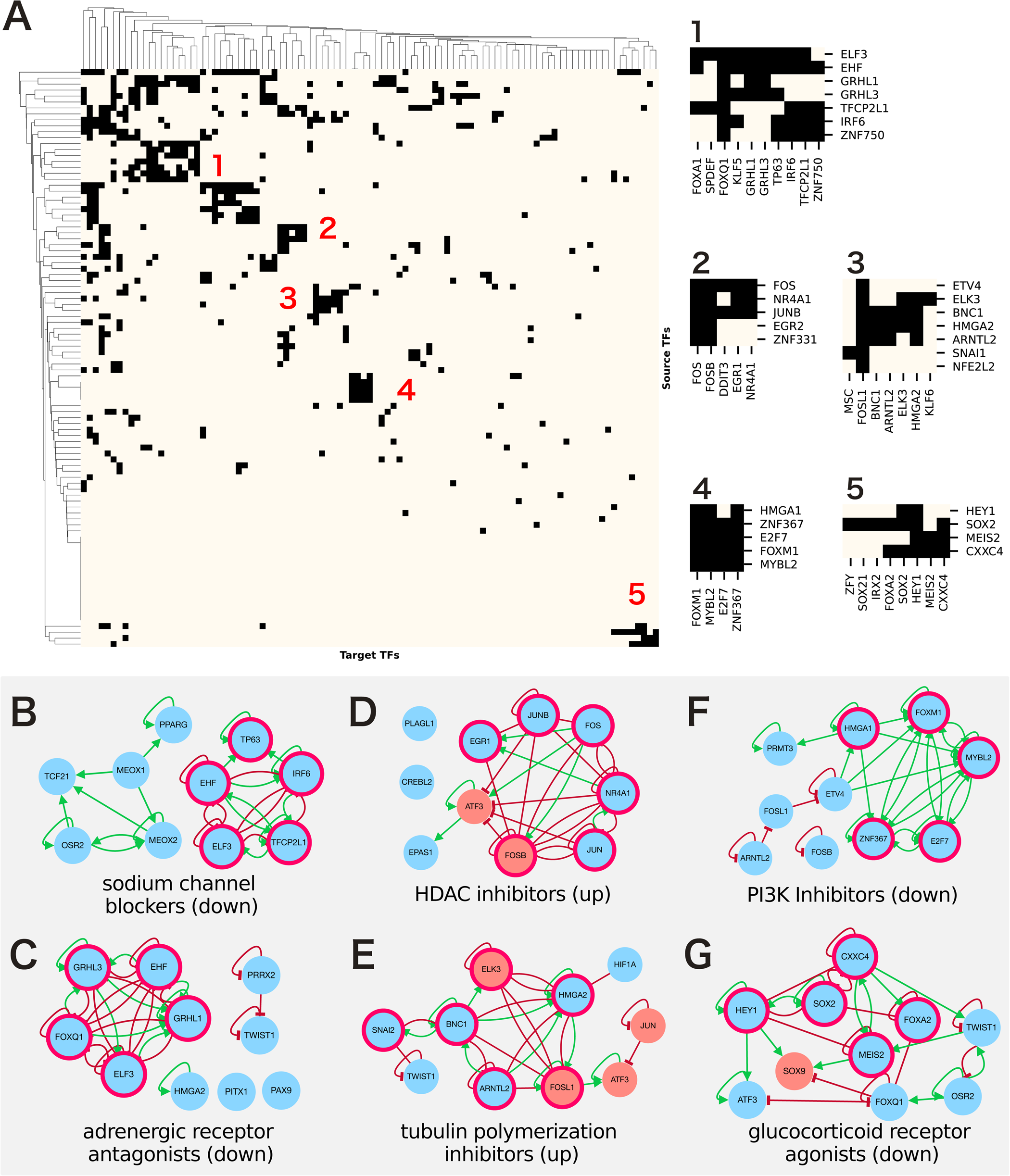
A: Hierarchically clustered binary adjacency matrix and selected co-regulating TF modules. B-F: Representative subnetworks for each identified module. Subnetwork TFs corresponding to module TFs are highlighted in pink. Module 1 consists of two MoAs and is represented by both B and C. Module 2 is represented by D. Module 3 is represented by E. Module 4 is represented by F. Module 5 is represented by G.

**Table 3.** Summary of each consensus gene set in the MoA Atlas, including the number of signatures used to compute each gene set, the number of unique drug perturbations included in the relevant signatures, and the final up and down gene set sizes.

| MoA | Signatures | Drugs, up | Up genes | Down genes |
| --- | --- | --- | --- | --- |
| dopamine receptor antagonist | 345 | 16 | 146 | 88 |
| cyclooxygenase inhibitor | 124 | 16 | 94 | 96 |
| adrenergic receptor antagonist | 77 | 16 | 104 | 116 |
| glucocorticoid receptor agonist | 136 | 15 | 153 | 177 |
| calcium channel blocker | 62 | 11 | 159 | 97 |
| histamine receptor antagonist | 54 | 11 | 126 | 116 |
| tubulin polymerization inhibitor | 193 | 10 | 231 | 189 |
| phosphodiesterase inhibitor | 65 | 10 | 113 | 129 |
| mTOR inhibitor | 703 | 9 | 146 | 135 |
| serotonin receptor antagonist | 45 | 9 | 181 | 148 |
| bacterial cell wall synthesis inhibitor | 25 | 9 | 191 | 167 |
| estrogen receptor agonist | 131 | 7 | 85 | 114 |
| adrenergic receptor agonist | 39 | 7 | 214 | 159 |
| PARP inhibitor | 107 | 6 | 83 | 135 |
| acetylcholine receptor antagonist | 23 | 6 | 279 | 313 |
| selective serotonin reuptake inhibitor (SSRI) | 69 | 6 | 211 | 128 |
| HDAC inhibitor | 1757 | 6 | 321 | 276 |
| sodium channel blocker | 47 | 5 | 206 | 150 |
| bacterial DNA gyrase inhibitor | 28 | 5 | 183 | 244 |
| HMGCR inhibitor | 84 | 5 | 166 | 144 |
| PI3K inhibitor | 727 | 5 | 160 | 106 |
| potassium channel blocker | 41 | 5 | 245 | 155 |
| ATPase inhibitor | 27 | 5 | 250 | 198 |
| HIV protease inhibitor | 23 | 5 | 315 | 257 |
| adenosine receptor agonist | 21 | 5 | 229 | 263 |
| AKT inhibitor | 78 | 4 | 180 | 95 |
| bacterial 50S ribosomal subunit inhibitor | 22 | 4 | 248 | 281 |
| DNA synthesis inhibitor | 38 | 4 | 195 | 190 |
| potassium channel activator | 15 | 4 | 267 | 309 |

#### Cluster 1: Adrenergic Receptor Antagonists (down)

The subnetwork representing Cluster 1 is composed of TFs enriched for genes downregulated by sodium channel blockers (TP63, IRF6, EHF, ELF3, TFCP2L1) and adrenergic receptor antagonists (GRHL3, EHF, GRHL1, FOXQ1, ELF3) (Figure 8B, 8C). These overlapping modules are generally associated with epithelial-associated TFs [104–107]. The TFs enriched in response to adrenergic receptor antagonists are associated with tumour suppressive TFs known to control the epithelial-mesenchymal transition (EMT). To test the role of these TFs in EMT regulation, 215 genes from an EMT-specific signature constructed via antibody-based profiling of 736 cancer cell lines [108] were submitted to ChEA3 [10] to identify likely TF regulators. All five TFs in the adrenergic receptor antagonist module were ranked within the top 12 most likely regulators of the EMT-specific genes (Supplementary Table S6). Beta-adrenergic signalling has been linked to the progression of cancer, in part through activation of the EMT [109]. Beta-blockers, a type of adrenergic receptor antagonist, have thus been explored as a potential cancer therapeutics, particularly triple-negative breast cancer [110, 111]. This suggests that one mechanism by which beta-blockers can produce an anti-cancer response is through activation of the tumour-suppressive, EMT-regulating known module of GRHL3/EHF/GRHL1/FOXQ1/ELF3.

#### Cluster 2: HDAC Inhibitors (up)

The subnetwork representing Cluster 2 includes six TFs from the subnetwork enriched for genes upregulated by HDAC inhibitors (EGR1, JUNB, FOS, NR4A1, JUN, FOSB) (Figure 8D). Four of these TFs, namely FOS, FOSB, JUN, and JUNB, are considered immediate early genes (IEG) [112]. IEGs are characterized by having low basal expression levels and rapid upregulation following extracellular stimuli. These genes are tightly regulated, in part because of their global impact on transcription. Their enrichment downstream of HDAC inhibitors suggests that IEGs the TFs and their targets are upregulated in response to HDAC inhibitors. This was confirmed by a study that found that administration of HDAC inhibitors in aging mice increases levels of fos-positive cells [113]. This suggests a possible mechanism by which HDAC inhibitors upregulate IEGs by reversing histone hypoacetylation at their promoter.

#### Cluster 4: PI3K Inhibitors (down)

The TFs enriched for genes downregulated by PI3K inhibitors and captured by Cluster 4 – HMGA1, FOXM1, MYBL2, ZNF367, and E2F7 – form a highly connected positive feedback module. These TFs are implicated in cell cycle control and cell proliferation [114–117]. Additionally, four of the five TFs, all except ZNF367, have been associated with the PI3K/AKT/mTOR signalling pathway [118–121]. This suggests that PI3K inhibitors suppress expression of TFs that belong to this subnetwork of nested positive feedback loops. ZNF367 is an understudied TF that has previously been tied to the progression of colon cancer via YAP1 signalling [122]. Its involvement in this tightly connected subnetwork module downstream of the PI3K signalling pathway is highly likely but needs to be confirmed experimentally (Figure 8F).

#### Cluster 5: Glucocorticoid Receptor Agonists (down)

The genes downregulated by glucocorticoid receptor agonists are enriched for targets of TFs tied to developmental programming related to the Wnt cell signalling pathway [123–125]. The gene encoding for the transcription factor HEY1 is a known to be upregulated by Notch signalling [126]. The TF CXXC4 is a known repressor of the Wnt pathway and a highly connected member within this module [123]. Glucocorticoid receptors have also been shown to repress Wnt signaling in human osteoblasts [127]. This suggests that glucocorticoid receptor agonists activate this module of TFs to suppress Wnt-related developmental genes that may have additional roles in normal physiology and disease (Figure 8G).

### The TF subnetworks of the aging atlas

The ChEA-KG aging atlas consists of aging-specific up and down gene sets for 24 tissues computed from the GTEx v8 publicly available RNA-seq dataset [128] (Supplementary Table S8). To explore how the various tissue-specific aging TF subnetworks may share components, a composite network was created by combining all edges and nodes from all the aging atlas subnetworks (Figure 9A). This network has visible clusters of TFs associated with specific tissues sharing similar aging specific regulatory modules (Figure 9B; Supplementary Figure S4). Clusters of co-regulating TFs were also identified by hierarchically clustering the adjacency matrix of the composite network (Supplementary Figure S5). One distinct network module was identified to regulate upregulated aging genes in breast, lung, blood vessel, thyroid, liver, and skin tissues with the TF Intestine-Specific Homeobox (ISX) appearing as a central node in this module (Figure 9C).

**Figure 9.**
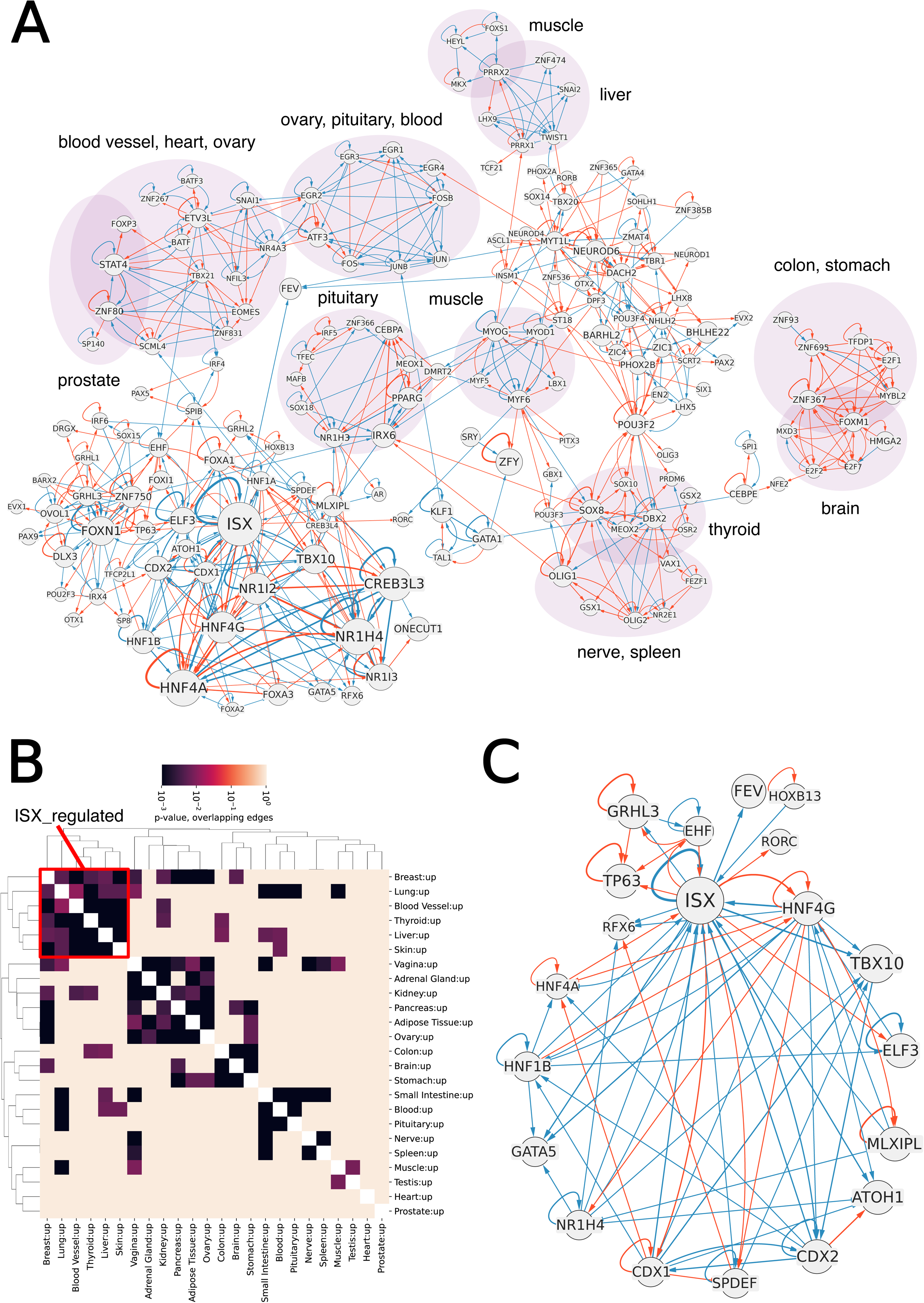
A: Cytoscape visualization of the Aging Atlas composite network. Node and edge sizes are proportional to the number of networks in which each appears. TF submodules corresponding to specific tissues are highlighted in purple and labeled. B: Heatmap of hierarchically clustered p-values representing similarity between regulatory subnetworks for genes upregulated in each tissue in response to aging. B: Subnetwork from (A) consisting of ISX and its immediate neighbors, based on gene sets upregulated in response to aging.

ISX is an intestinal tissue master regulator that is also responsible for controlling vitamin A production [129]. ISX has been also suggested as a potential prognostic and therapeutic target for hepatocellular carcinoma (HCC) [130]. The ISX associated subnetwork includes three hepatocyte nuclear factors (HNFs), namely HNF4ɑ, HNF4ɣ, and HNF4β. These TFs are primarily expressed in the liver, pancreas, and kidney and their dysregulation is implicated in diabetes [131]. Other TFs found in the subnetwork are also involved in liver and pancreatic functions including NR1H4, a regulator of bile acid synthesis [132]; MLXIPL, a master regulator of lipogenesis [133]; and RFX6, which regulates pancreatic islet cell differentiation [134]. The remaining TFs in the subnetwork, CDX1, CDX2, ELF3, GATA5, TBOX10, SPDEF, and AOTH1, are cellular differentiation regulators in both embryonic and adult tissues [69, 70, 135–137]. Dysregulation of developmental programs in adult tissues has recently been suggested as a cause of aging and age-related disease [138]. This module therefore implicates metabolic function and tissue development in the process of aging.

## DISCUSSION

The formation of the ChEA-KG GRN presents a unique new method to reconstruct in-silico gene regulatory networks. By combining a massive collection of diverse gene expression signatures with a comprehensive TF enrichment analysis, a reliable data driven GRN was constructed. The GRN contains almost all known human TFs, expanding on the size and coverage of most current GRN reconstruction techniques. For example, it includes almost all known human TFs (n=1,559) in its background GRN. In contrast, TRRUST [14], a highly used human GRN reconstructed from text mining, is considerably smaller (n=800). The accuracy of the GRN edges is supported by the benchmarking results. However, testing it against other sources of known TF-TF interactions would strengthen the confidence of the links. Ultimately, testing such regulatory interactions in-vitro and in-vivo would fully confirm the inferred regulatory relationship. This can be achieved by knockdown, knockout, and over-expression of the source TF followed by gene and protein expression of the target TF. Examining the topology of the GRN identified clear clusters with known shared functions for each module. For example, we identified a module of TFs that is related to the cell cycle, immune response, differentiation, and development.

The ChEA-KG web server application facilitates the querying of the GRN for knowledge discovery. The enrichment analysis feature of ChEA-KG can be used to uncover how the most enriched TF master regulators are forming subnetworks of tight regulatory interactions. In the future, such subnetworks could provide the basis for dynamical simulations to further understand how groups of TFs work together as a functional regulatory unit. In addition, the GRN can be expanded to include non-TF-encoding genes such as kinases, chromatin modifiers, and other relevant components that can enrich the contents of each subnetwork.

The several atlases of ChEA-KG demonstrate how the GRN can be used to systematically identify known master regulator subnetworks and propose new regulatory mechanisms in a variety of contexts. For example, the cell type specific subnetworks also contain TF unknown to be relevant to the specific cell types, potentially leading to new discoveries. This includes, for example, the role of MRFs in regulation of cardiac muscle cells, or the role of ELK3, an understudied member of the ETS family, in mesenchymal stromal cells regulation. The cancer specific subnetworks demonstrate how ChEA-KG can be integrated as part of a transcriptomics workflow to better understand the distinguishing factors between cancer subtypes. The MoA atlas subnetworks reveals connections between known regulatory modules and common drug MoAs. For example, a connection between HDAC inhibitors and IEGs is evident. Also, the connection between adrenergic receptor agonists and EMT regulators is interesting. Finally, the global network that is made of all the upregulated aging atlas tissue subnetworks reveals a possible new master regulatory module of aging, featuring metabolic and developmental TFs that are cantered around ISX.

## CONCLUSIONS

Overall, ChEA-KG presents several features not available in existing tools. In addition, the ChEA-KG network visualization interface enables users to easily see regulatory subnetworks that connect TFs with signed and directed links. Currently, we are not aware of a dynamic network visualization feature in any of the TF enrichment analysis servers/apps. Importantly, ChEA-KG has four atlases that visualize the master regulatory subnetworks for 131 human cell types, 69 tumour subtypes from 10 cancers, 30 drug MoA signatures, and 24 aging tissue signatures. By using the ChEA-KG GRN as a background regulatory network, ChEA-KG can provide insights to biologists about not only the TFs that are most relevant to their data, but also how these TFs form a regulatory module that works in concert to induce the observed changes in gene expression, adding more context to TF enrichment analysis results.

## LIST OF ABBREVIATIONS

TF: Transcription Factor
GRN: Gene regulatory network
ChIP-seq: Chromatin immunoprecipitation followed by sequencing
ChIP-chip: Chromatin immunoprecipitation followed by microarray
DNase-seq: DNAse sequencing
FAIRE-seq: Formaldehyde-assisted identification of regulatory elements followed by sequencing
ChIA-PET: Chromatin interaction analysis with paired end tag sequencing
PWM: Positional weight matrices
TFEA: Transcription factor enrichment analysis
KG: Knowledge graph
TSS: Target set swap
ND: Node draw
UMAP: Uniform manifold approximation projection
TF-IDF: Term frequency-inverse document frequency
KG-UI: Knowledge Graph User Interface
UI: User interface
MoA: Mechanism of Action
LINCS: Library of Integrated Network-based Cellular Signatures
L1000 FWD: LINCS L1000 Fireworks Display
GTEx: Genotype-Tissue Expression Project
CPTAC3: Clinical Proteomic Tumor Atlas Consortium
GMT: Gene matrix transpose
MSCs: Mesenchymal stromal cells
MRFs: Myogenic regulatory factors
EMT: Epithelial-to-mesenchymal transition
EnMT: Endo-cardial mesenchymal transition
NIH: National Institutes of Health
LSCC: Lung squamous cell carcinoma

## AVAILABILITY OF DATA AND MATERIALS

The ChEA-KG web-server application is available at: https://chea-kg.maayanlab.cloud/. The node and edge lists for the ND-filtered, TSS-filtered, and unfiltered GRNs, as well as the marker gene sets for both the Cell Atlas and Cancer Atlas, are available on the ChEA-KG site, https://chea-kg.maayanlab.cloud/download_files. The ChEA-KG source code is available from GitHub at: https://github.com/MaayanLab/ChEA-KG-UI.

<u>Project name:</u> ChEA-KG

<u>Project home page:</u> https://chea-kg.maayanlab.cloud/

<u>Operating systems</u>: Platform independent

<u>Programming language</u>: Python, TypeScript, React

<u>License:</u> GPL 3.0

## FUNDING

This work was supported by the National Institutes of Health [OT2OD036435, U24CA264250, OT2OD030160, U24CA271114]. Funding for open access charge: National Institutes of Health [OT2OD036435].

## AUTHOR CONTRIBUTIONS

AB and JEE: data curation, formal analysis, software, validation, visualization, writing – original draft; AL: methodology, software; HYC: initial data curation, formal analysis, software; SJ: supervision, project administration; AM: conceptualization, funding acquisition, investigation, methodology, supervision, project administration, writing – original draft, writing – review & editing.

---

**Fig. S1.**
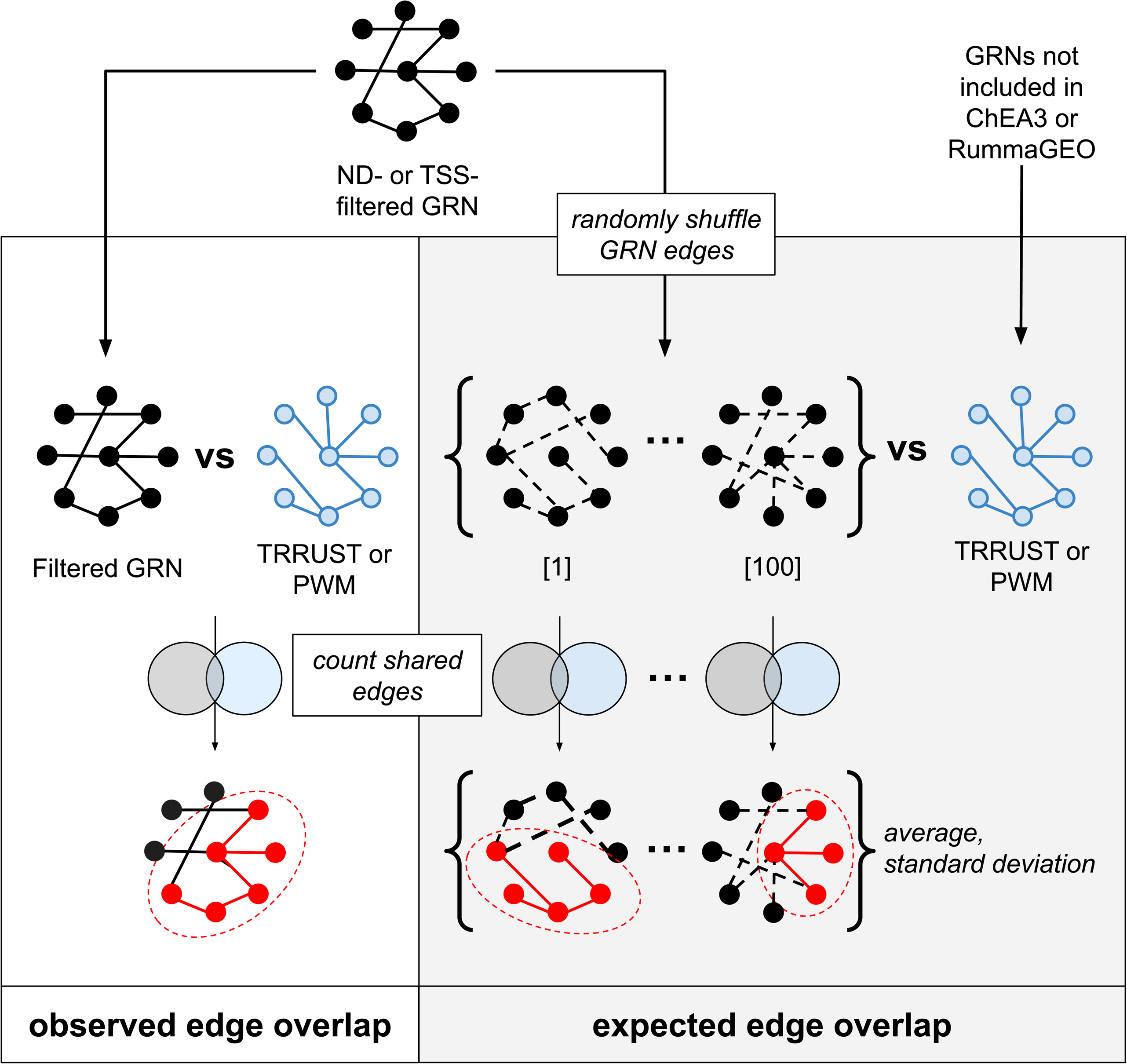

**Fig. S2.**
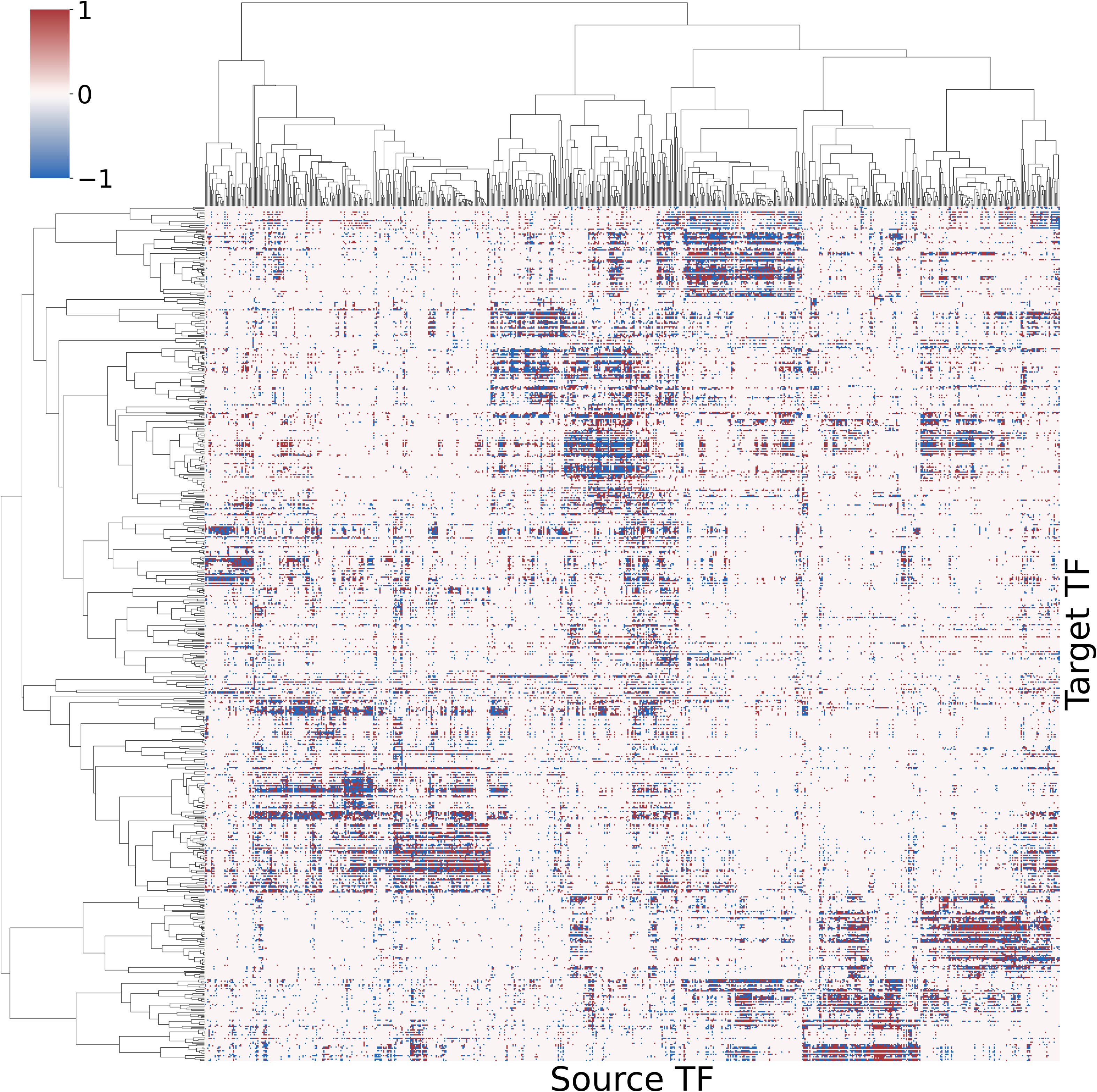

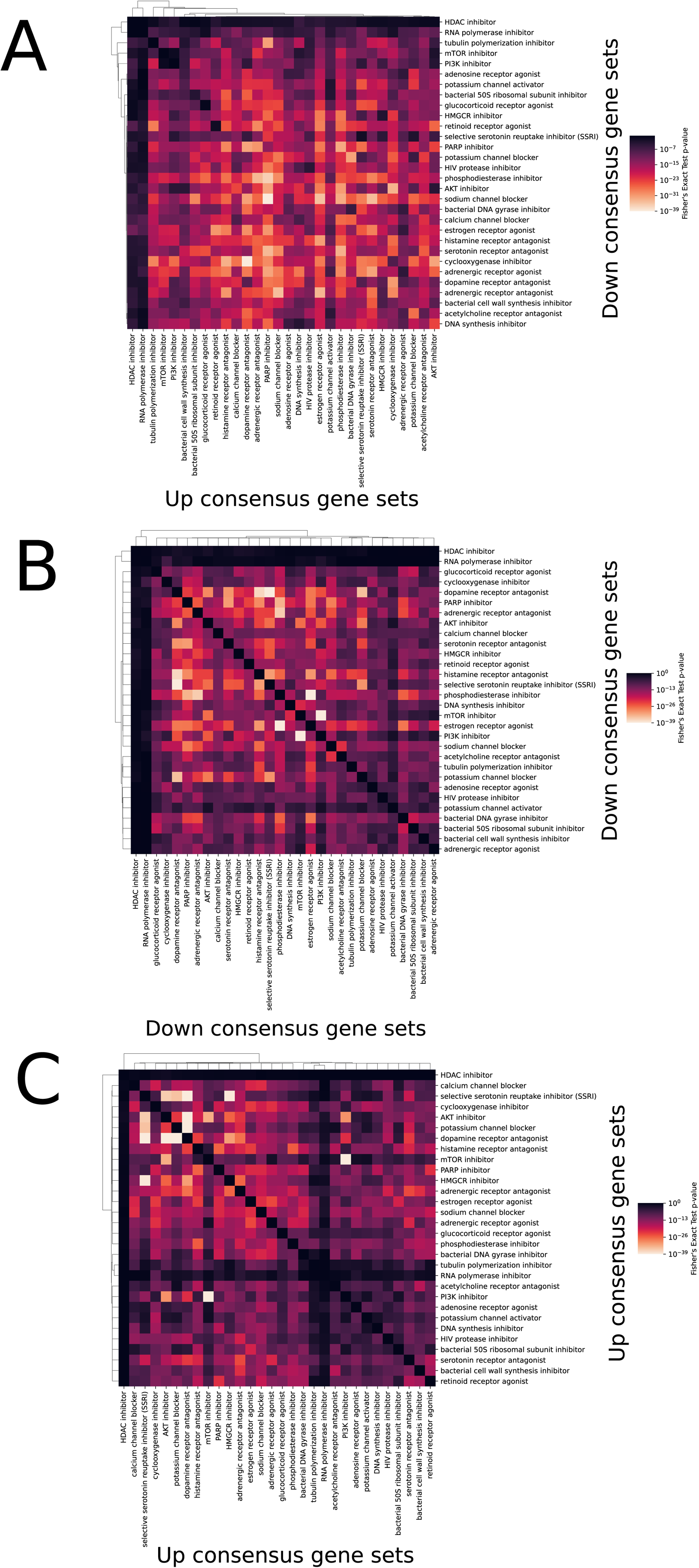

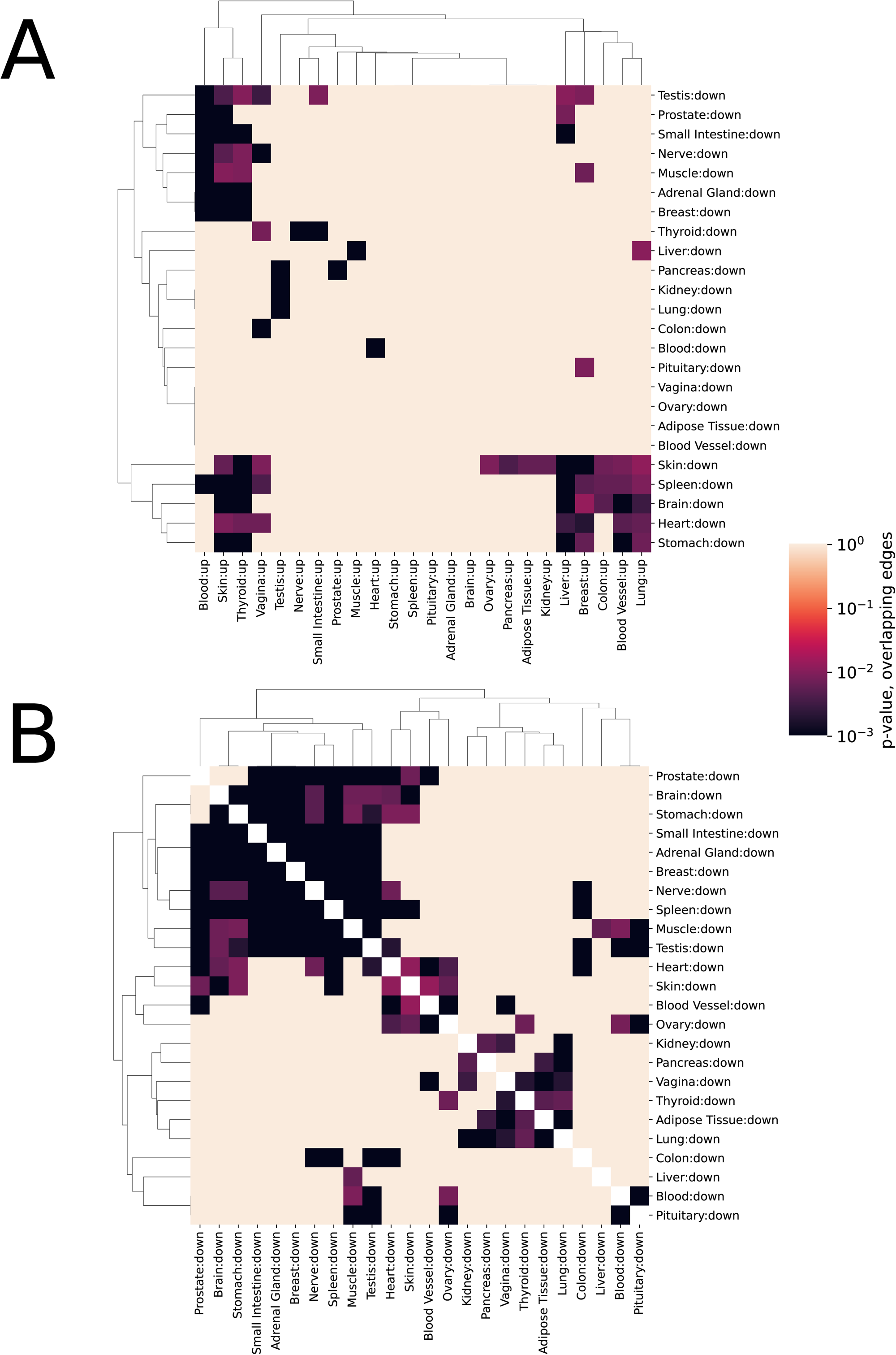

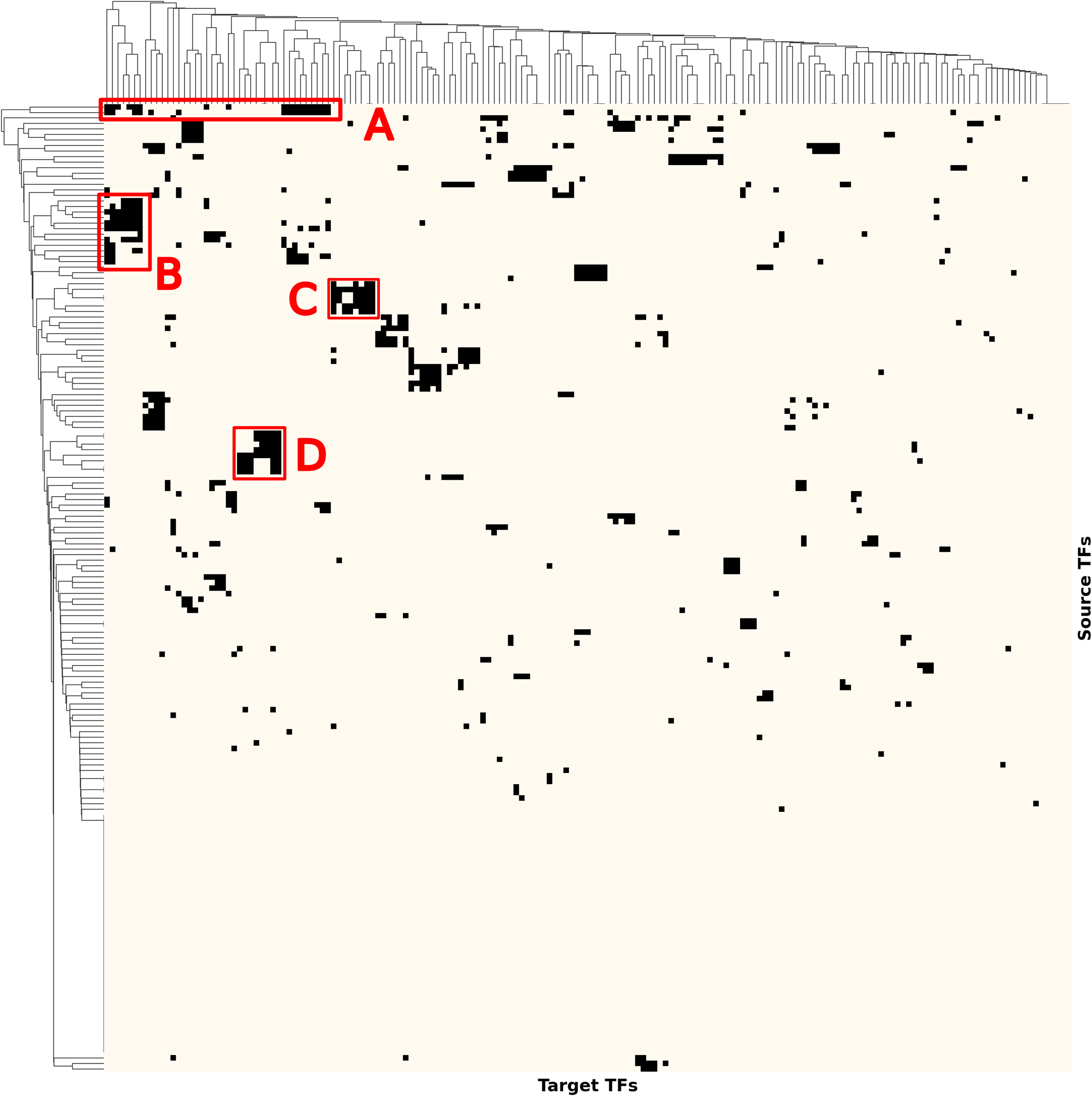

## Notes

https://chea-kg.maayanlab.cloud

### Competing Interest Statement

The authors have declared no competing interest.

### Summary of Updates

This version of the manuscript includes the implementation of several additional atlases to the ChEA-KG application including: - A cell types atlas created from gene sets extracted from CellMarker, ASCT+B, PanglaoDB, Tabula Sapiens, Descartes, Azimuth, and TISSUES - A pan-cancer atlas for subtypes of 10 tumor types extracted from CPTAC An aging atlas across human tissues using the GTEx transcriptomics data - Mechanisms of action for drugs atlas using the gene expression data from the LINCS L1000 dataset

